# GraphComm: A Graph-based Deep Learning Method to Predict Cell-Cell Communication in single-cell RNAseq data

**DOI:** 10.1101/2023.04.26.538432

**Authors:** Emily So, Sikander Hayat, Sisira Kadambat Nair, Bo Wang, Benjamin Haibe-Kains

## Abstract

Interactions between cells coordinate various functions across cell-types in health and disease states. Novel single-cell techniques enable deep investigation of cellular crosstalk at single-cell resolution. Cell-cell communication (CCC) is mediated by underlying gene-gene networks, however most current methods are unable to account for complex interactions within the cell as well as incorporate the effect of pathway and protein complexes on interactions. This results in the inability to infer overarching signalling patterns within a dataset as well as limit the ability to successfully explore other data types such as spatial cell dimension. Therefore, to represent transcriptomic data as intricate networks, complementing gene expression with information from cells to ligands and receptors for relevant cell-cell communication inference, we present GraphComm - a new graph-based deep learning method for predicting cell-cell communication in single-cell RNAseq datasets. GraphComm improves CCC inference by capturing detailed information such as cell location and intracellular signalling patterns from a database of more than 30,000 protein interaction pairs. With this framework, GraphComm is able to predict biologically relevant results in datasets previously validated for CCC, datasets that have undergone chemical or genetic perturbations and datasets with spatial cell information.

## Introduction

Understanding multicellular organisms and their ability to function requires an in-depth understanding of cellular activities^1,2^. This can be achieved by studying the signalling events that will induce responses and downstream effects. The most commonly studied signalling events typically involve the binding of a secreted ligand to a cognate receptor, either in an intercellular (paracrine)^3^ or intracellular (autocrine)^4^ fashion. The importance of these ligand-receptor interactions in biology has led to an increasing interest in computationally uncovering patterns of cell-cell communication (CCC)^5^ and resulting phenotypic effects in healthy, perturbed, and disease conditions. Subsequently, changes in CCC will be helpful in improving our understanding of tissue function^5^ and disease progression^5,6^. Direct cell-cell communication has been shown to be crucial for tumour progression, which enables the transfer of cellular cargo from non-cancerous to cancerous cells^7,8^. Robust computational tools to model this type of cell-cell communication in tumours can also aid in identifying key patient-selection biomarkers^9^ and improving therapeutic approaches for drug response^10^.

One of the major challenges in determining CCC is the lack of ground truth in defining potential ligand-receptor pairs, which limits the subsequent validation^11^. Identifying interactions in their native microenvironments often requires expensive experiments or extensive domain knowledge in detecting active interactions^12^. To allow for a more accessible method for studying CCC, single-cell transcriptomics has been incorporated in utilising evidence of gene expression to infer CCC activity. As single-cell RNAseq is increasingly used to study cell types and states, there is an imminent need for computational methods that can perform prediction of validated ligand-receptor activity with count matrices of gene expression. Methods such as CellPhoneDB^13^, Crosstalkr^14^, Connectome^15^, NicheNet^16^ and CellChat^17^, have paved the way for developing CCC methods that can generate results at both the bulk and single-cell level. Based on the results from these methods, there has also been increasing use in applying CCC methods to different modalities such as spatial transcriptomics data^18–20^, where cell coordinates could provide a basis for visualising CCC predictions with respect to spatial adjacency.

There are, however, multiple limitations that must be addressed to improve the accuracy and biological relevance of CCC prediction. There is a dearth of a consistent ground truth for validated ligand-receptor (LR) pairs with accompanying annotation, such as protein complex information and pathway information^11^. In an effort to maximise availability of information during the prediction process, there are opportunities to identify new methods for representation methods for LR interactions and properties. Deep learning is a suitable candidate for application, proven from its performance in other applications of biological networks^21^. Using a more detailed view of a ligand-receptor ground truth through a deep learning model can allow for results of CCC activity reflective of a change in transcriptomic values.For example, LR information can be used in conjunction with context-specific transcriptomic data as the expression strength of ligand and receptor-encoding genes has been previously directly linked to their interaction probability^5^. By introducing the utilisation of deep learning in a CCC prediction framework, novel ligand-receptor interactions can be identified that are indicative of a new cell role or cell function in different contexts.

To address these issues, we present GraphComm, a new Graph-based deep learning method to predict CCC from single-cell RNAseq data. GraphComm uses more detailed labels of ligand-receptor annotation (such as protein complex and pathway information) as well as expression values and intracellular signalling to construct cell interaction networks using feed-forward mechanisms to learn an optimal data representation and predict probability of CCC interactions. GraphComm is able to compute communication probabilities that represent relationships between cell groups, ligands and receptors. Therefore, for any given ligand-receptor link, a communication probability based on computed embeddings can be extracted and used to rank CCC activity. GraphComm’s designed architecture allows for rich information regarding both cellular signalling networks and transcriptomic information to be captured in predictions and enables GraphComm to prioritise multiple interactions at once. Using single-cell transcriptomics data from embryonic mouse brain, human hearts and cancer cell lines, we demonstrate the utility of GraphComm in aligning with previously identified ligands and receptors, identifying biologically relevant changes in CCC in cancer cells after drug perturbation, and predicting cell-cell interactions in spatial microenvironments. Therefore, GraphComm has the potential to be a robust and translatable computational framework that can uncover small and large-scale communication patterns in transcriptomic data.

## Results

### GraphComm overview

Using a curated ligand-receptor database and single-cell transcriptomic data, GraphComm utilises directed graph representations of cell-gene networks to infer meaningful, validated and novel ligand-receptor interactions between two cell types or cell clusters **(Figure 1)**. The prediction pipeline of GraphComm is split into two distinct steps: 1) feature representation learning using a prior model **(Figure 1A),** and 2) using transcriptomic information to predict cell-cell communication present in a single-cell dataset **(Figure 1B)**.

**Figure 1:**
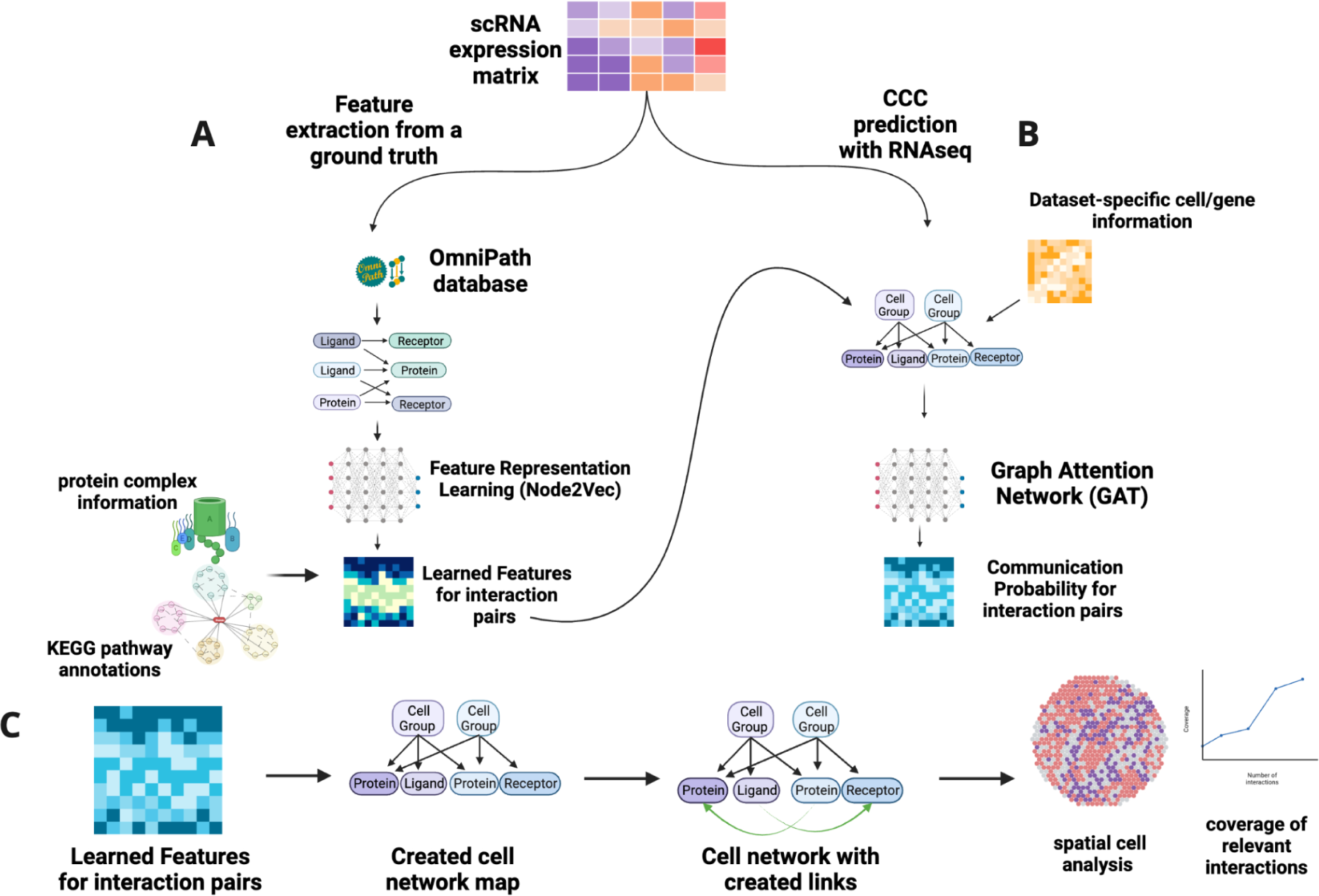
Schematic outlining the architecture of GraphComm to make CCC predictions from scRNAseq data **(A)** GraphComm utilises a scRNAseq dataset with a curated ligand-receptor database to construct a directed graph reflective of the CCC ground truth. Feature Representation is implemented to extract positional information for ligands and receptors within the directed graph and scaled accordingly with protein complex and pathway information **(B)** GraphComm constructs a second directed graph representing the relationship between cell groups and source/target proteins. Annotating this second directed graph with transcriptomic information, cell group information and positional features from the Feature Representation learning step to receive updated numerical node features via a Graph Attention Network. Node Features can then be used via inner product to compute communication probability for all possible ligand receptor pairs. **(C)** Computed communication probabilities via the Graph Attention Network can be combined with the second directed graph to construct ligand-receptor links with top-ranked CCC activity, which can be used for visualisation of activity at the ligand-receptor and cell group level. **Created in** https://BioRender.com

### Feature Representation Learning using a Prior Model

Utilising features for validated ligand-receptor pairs from ground truth is critical for inferring CCC in transcriptomic data to prioritise cellular processes such as co-expression. This requires a comprehensive, large curated ligand-receptor database, for which we currently use OmniPath^22^. In addition to providing more than 30,000 validated intracellular interactions and more than 3,000 validated intercellular interactions, OmniPath is accompanied by protein complex and pathway information^22,23,24^. Incorporating this information has proven to be successful previously in methods such as LIANA^11^ and CellChat^17^, incorporating factors on gene expression from protein sub-unit/pathway and extracting more complex patterns from CCC. First, GraphComm utilises an input single-cell RNAseq expression matrix and identifies all significantly expressed ligands, receptors and intracellular proteins present in the dataset. A directed graph using these proteins is constructed, with edges drawn from source to target only if the link occurs with validation in the OmniPath Database. Quantitatively, the graph will have (*# of source and target proteins present in the dataset)* nodes *(# of validated source-target protein interactions)* edges. Using this updated directed graph, Feature Representation Learning is conducted via the Node2Vec^25^ framework, which will compute new positional numerical embeddings for each node present in the graph by utilising a loss function that samples false connections during training, forcing the model to focus on true relationships. Once Representation Learning is completed, the processed outputs of this task are then scaled with a separately computed numerical matrix of shape *(# of ligands) x (# of receptors)*, containing numerical values detailing ligand and receptors’ correlation from the ∼8,022 protein complexes and ∼7500 KEGG pathways members (see Methods) present in the OmniPath Database. These scaled constructed features are used as downstream input features for validated ligand-receptor links that can be used to further predict CCC activity, where larger values are assigned to ligand-receptor pairs that are co-expressed in the same pathways or in the same protein complexes and favouring their pairing.**(Figure 1A).**

### scRNAseq infers cell-to-cell communication probability

After numerical features have been extracted using context-independent information from the OmniPath Database, communication probability is calculated for all possible source/target protein links, both intracellular and intercellular. This calculation begins with the input of a given scRNAseq dataset by identifying defined cell groups within the dataset (such as cell types obtained after Louvain/Leiden^26^ clustering) and each group’s differentially expressed proteins. A new directed graph is constructed using nodes of three types: cell groups/clusters, source proteins and target proteins. Edges are drawn from a given cell group to a source/target protein if the expression of the source/target protein in the cell group is significantly up-regulated. This newly created graph resembling the potential cellular network is further annotated by a combination of previously obtained positional embeddings and new contextual information (see Methods). The annotated directed graph is fed as input to a Graph Attention Network (GAT)^27^ for 100 training epochs. Computed initial biological node embeddings for all nodes are continuously updated, skewing towards a binary ground truth favouring interacting intracellular ligands and receptors (which are interactions defining the basis of cell communication aside from other PPI). Finally, a table containing computed interaction probability for all possible source/target protein links (the total number of possible links dependent on the number of protein-coding genes in the dataset) are obtained via inner product computing of the GAT output, which are used as communication probability **(Figure 1B).**

### Cell Communication results and visualisation

To visualise possible ligand-receptor links in a dataset, the new matrix from the last step of the GraphComm framework can be utilised in combination with the input graph **(Figure 1C)**. For each possible ligand/receptor pair in the original input graph, a new link is created with a corresponding computational probability. This allows for ranking of ligand-receptor pairs for inference and visualisation, as well as identifying source and destination cell groups. Additionally, using the chosen datasets GraphComm can perform analyses to confirm previously validated activity and report the potential of new communication (**Figure 2).**

**Figure 2:**
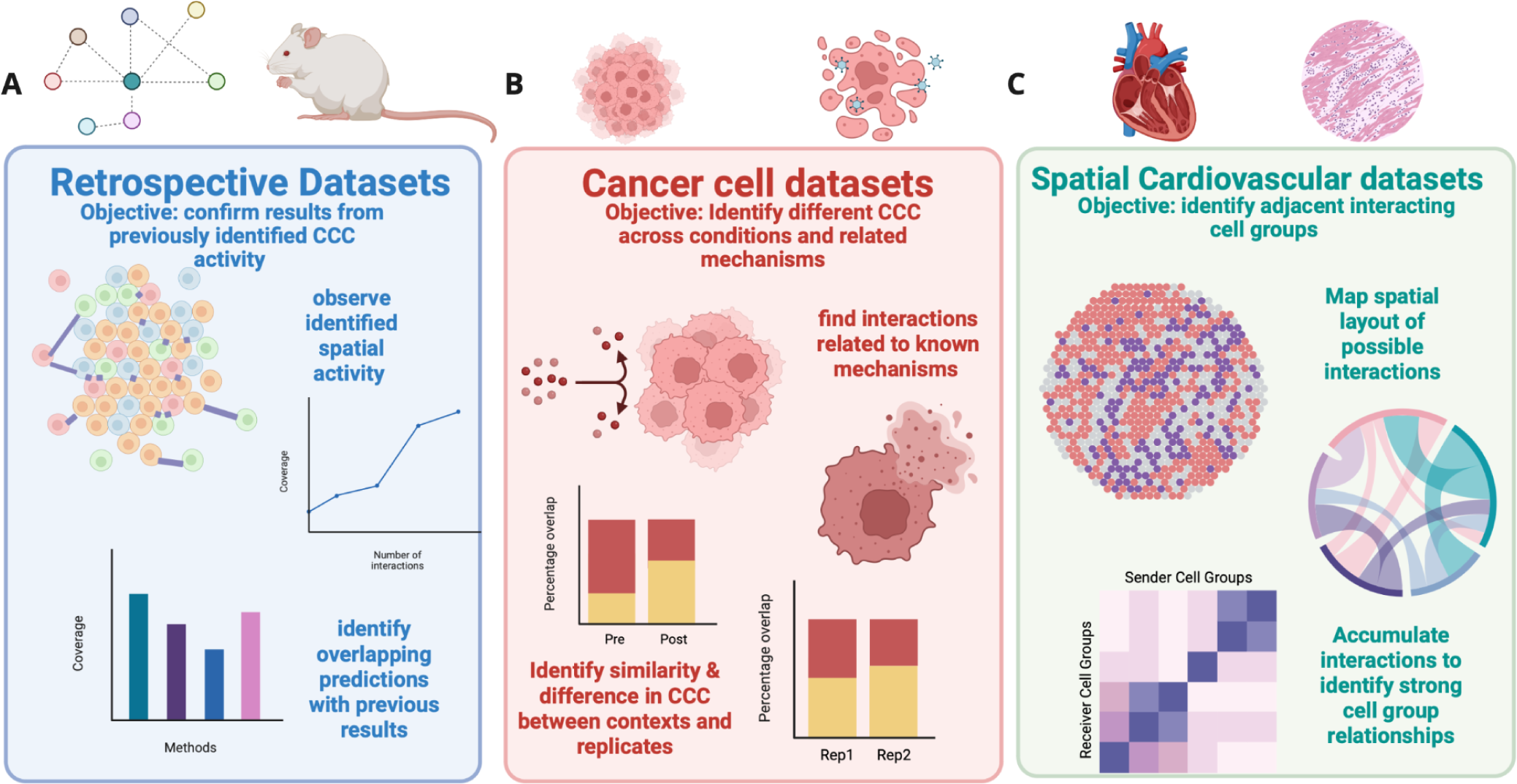
Schematic of the validated datasets that GraphComm performs inference on and the purpose for their respective benchmarks. **(A)** Datasets with validated CCC activity, from embryonic mouse brain, were used in GraphComm prediction to confirm predictions aligned with the previously validated results. Measures of success in those benchmarks include the plotting of spatial activity and top-ranked interactions to compare with the ground truth.**(B)** Datasets of treated cancer cells from PC9 cell lines were used in GraphComm prediction to identify how our framework can identify different CCC across conditions and connect communication activity to related, well-studied mechanisms. Benchmarks will compare the similarity and difference in interaction overlap between datasets of different conditions (pre and post treatment) and biological replicates. Furthermore, validity in predicted interactions can be strengthened by highlighting ligands and receptors that have been previously studied in cancer cell drug resistance and sensitivity mechanisms. **(C)** Datasets containing spatial transcriptomics, from human hearts following myocardial infarction, were used in GraphComm prediction to identify potentially interacting adjacent cell groups. With location information within an individual slide, the likelihood of an interaction can be correlated to its predicted communication probability. By also identifying the number of interactions in one zone across all available slides, crosstalk between two cell groups can be analysed for their relevance towards the dataset. **Created in** https://BioRender.com

### GraphComm predicts cell-cell communication in retrospective datasets

To assess the ability of GraphComm to identify previously validated inter-cell group communications, we applied our method on an embryonic mouse brain published by Sheikh et. al^28^. In the original study, the authors identified patterns of communication between cell groups, including 1,710 unique ligand-receptor interactions that were all validated. The results found in this publication were able to reveal to the author underlying mechanisms and patterns that are critical in mouse brain development and embryogenesis.

We used GraphComm to infer, among the top-ranked interactions of the prediction set, how many were also present as important interactions in the original publication **(Figure 3A)**. To evaluate the robustness of GraphComm, we first conducted 100 randomization trials, in which results of GraphComm were computed, trained on a randomised version of the ground truth. It was found that across 100 randomised iterations, on average about 45% of the top 100 interactions would contain a ligand or receptor present in the previous publication set. In contrast, it seemed that a true inference from GraphComm, using the full correct ground truth, was able to prioritise important ligands and receptors more accurately achieving a range of 48-55% of the top 100 interactions containing a previously seen ligand or receptor. **(Figure 3B).** The overlap with the publication of GraphComm’s top 100 interactions is either comparable or slightly lower than other methods (averaging across all interactions, a coverage of 1-3% less) **(Extended Data Figure 1)**. The difference in performance can be connected to the difference in prioritisation of certain interactions, as GraphComm uses a larger pool of interaction candidates than other methods and therefore has a higher probability of detecting interactions with ligands and receptors not previously found). Therefore, GraphComm was able to identify previously validated ligand and receptor pairs and could possibly lead to the highlighting of novel activity within the embryonic mouse brain.

**Figure 3:**
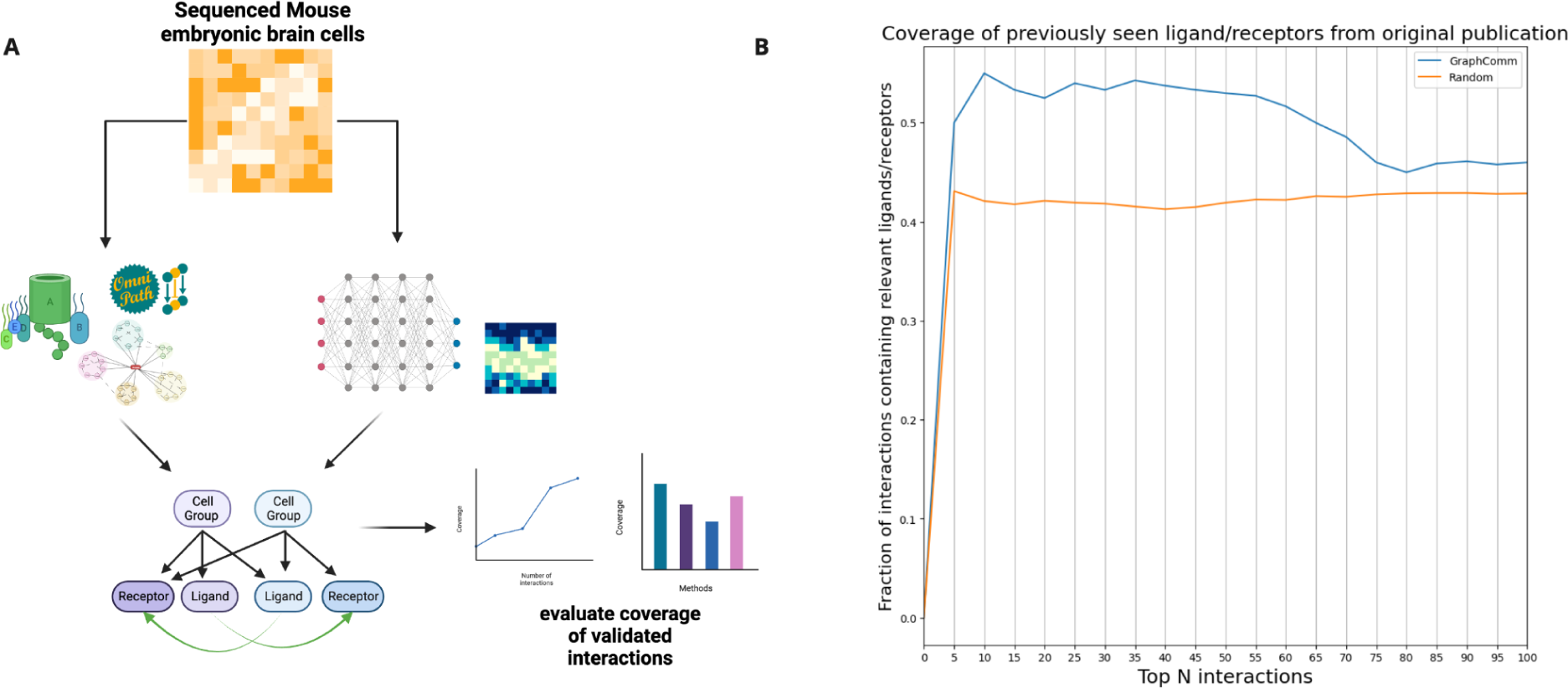
**(A)** GraphComm can identify interactions previously validated *in silico* in an embryonic mouse brain **(B)** In comparison to 100 randomised iterations, GraphComm can identify more interactions containing previously validated ligands and receptor in the top 100 interactions with ligand and receptor candidates **Created in** https://BioRender.com

### GraphComm predicts changes in cell-cell communication due to genetic and chemical perturbations

To understand the potential of GraphComm in finding pathways and CCC affected by drug treatment, network inference was conducted on scRNA cancer cell lines pre- and post-drug treatment. The dataset consists of a pre-post treatment set of PC9 lung adenocarcinoma cell lines^29^, with cell populations at Day 0, 3, 7 and 14 obtained after being treated with Osimertinib (a tyrosine Kinase inhibitor^30^). We used GraphComm to (*i*) identify the condition-specific CCC and (*ii*) identify more common interactions in two biological replicates than two datasets of different conditions.In this task, the ground truth is still limited to the information we had available in our previous benchmarks – validated Ligand-Receptor (LR) interactions in the OmniPath database and known pathway information annotated via KEGG^23^. To identify how GraphComm provides inference on single-cell datasets of different and the same conditions, two sequencing timepoints from a cancer cell line dataset were used. One dataset was sequenced before treatment (day 0) and two biological replicate datasets were sequenced 7 days after treatment **(Figure 4A)**.

**Figure 4:**
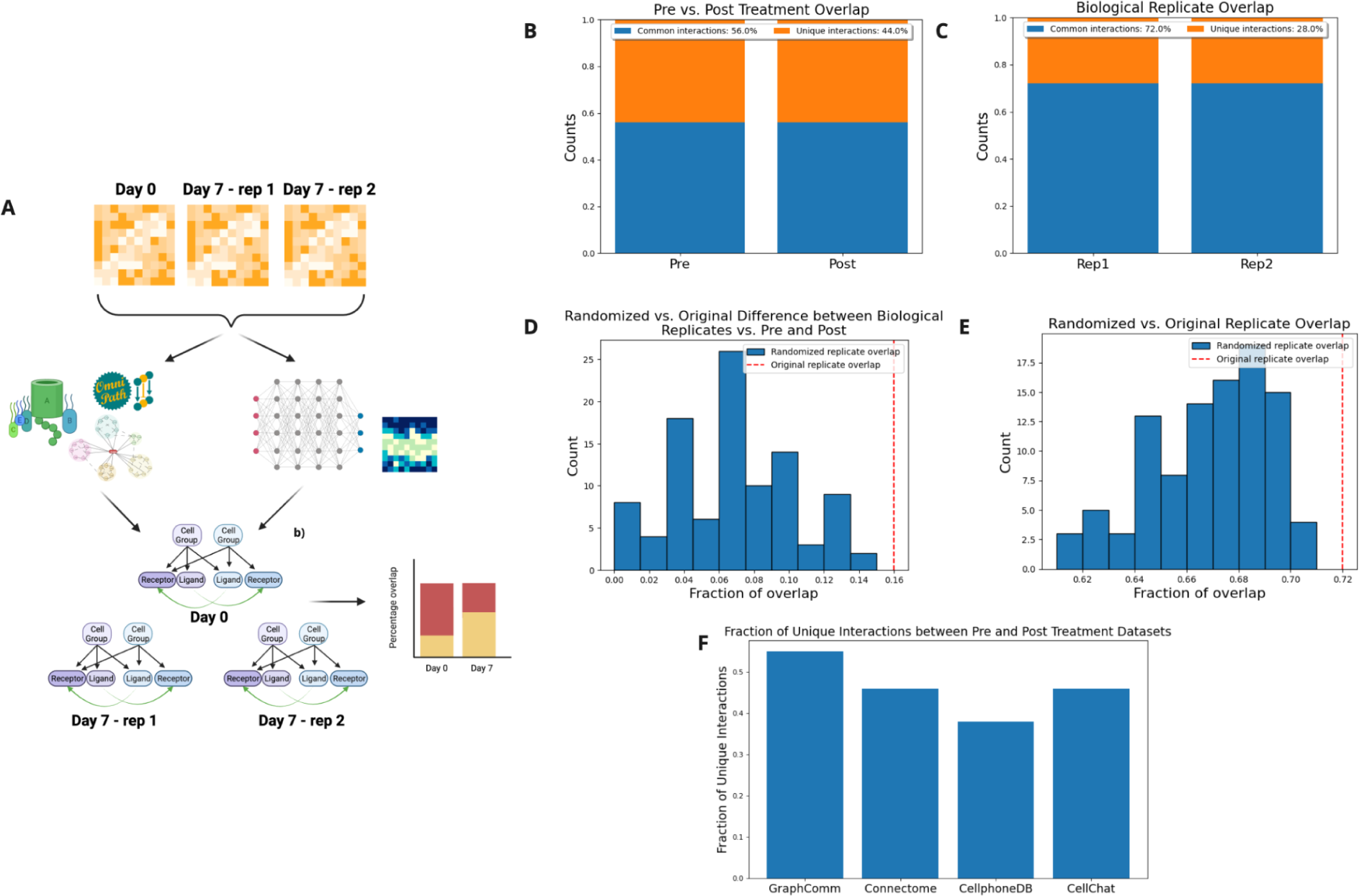
**(A)** GraphComm can identify differences in interactions among datasets that have the potential to be indicative of change in condition. Using multiple datasets of PC9 lung adenocarcinoma cell lines sequenced at different timepoints before and after treatment with osimertinib including two biological replicates, the fraction of overlapping interactions can be compared. This allows for the interpretation of how GraphComm can i) identify more unique interactions in two datasets of different condition than two biological replicates and ii) identify more common interactions in two biological replicates than two datasets of different condition **(B)** Between one PC9 dataset at day 0 (pre-treatment) and one PC9 dataset at day 7 (post-treatment), GraphComm is able to prioritise unique top-ranked interactions, with an overlap of 56% **(C)** In comparison to overlap between pre and post treatment datasets, GraphComm achieves a greater overlap between two day 7 (post treatment) biological replicates, with an overlap of 72% **(D)** In comparison to top-ranked interactions of 100 randomised iterations, GraphComm achieves a larger overlap difference of biological replicates vs pre and post treatment datasets using the original directed graph and ground truth **(E)** In comparison to top-ranked interactions of 100 randomised iterations, GraphComm achieves significantly better overlap between two biological replicates using the original directed graph and ground truth **(F)** In comparison to other CCC methods, GraphComm achieves a higher number of unique interactions between pre and post treatment datasets, potentially indicating GraphComm performs better at identifying interactions that can be indicative of a change in condition.**Created in** https://BioRender.com

We first looked at the top 100 validated intercell interactions in all 3 datasets to identify overlap between biological replicates and different conditions. In this benchmark, the ideal result would be quantitatively observed in a close to 0% overlap between pre- and post-treatment interactions and a close to 100% overlap between post-treatment biological replicate interactions. In this assessment, it was observed there was a 72% overlap between the two post-treatment biological replicates. The number of common interactions between biological replicates was larger than the 56% overlap between pre and post treatment datasets **(Figure 4B,C)**. To test the significance of the overlap between biological replicates versus pre- and post-treatment datasets, we conducted 100 randomised iterations of the analysed datasets and were unable to find the same difference in overlap as the original test set (p < 0.01) (**Figure 4D)**. This difference in common interactions pre- and post-treatment allows for the discovery of new interactions within the respective datasets that can lead to new findings in PC9 sensitivity or chemoresistance^31^. To once again evaluate robustness of GraphComm’s prediction capabilities, we conducted 100 randomised inferences using a version of GraphComm trained on a randomised ground truth on both post treatment datasets. The number of common interactions predicted between post-treatment datasets was also significantly higher in the original inference results in comparison to all results from the randomisation trials (p < 0.01). (**Figure 4E)**. In comparison to results from other established CCC methods, GraphComm outperformed other methods in identifying top validated intercellular interactions between pre and post treatment than other methods. (**Figure 4F).** Therefore, the results from GraphComm imply that our framework identifies, more than other methods, unique and biologically robust interactions in pre- and post-drug treatment cancer datasets.

### GraphComm predicts cell-cell communications in spatially adjacent cells

To assess GraphComm’s breadth of discovering probable CCC, its ability to predict interactions in accordance with other modalities can be assessed. For this inference benchmark, GraphComm is applied on spatial transcriptomics data obtained from human hearts post-myocardial infarction (MI), where different regions of the impacted hearts (Ischemic, remote, border and fibrotic zones) affected by myocardial infarction and control^32^ (unaffected) were sequenced.For all regions of the heart sequenced, cell expression is also accompanied by spatial coordinates which allows for both cell and gene to be quantitatively weighted. The overall objective of this benchmark is to demonstrate GraphComm’s ability to learn graph structure in agreement with its spatial microenvironment, i.e., cell positioning and adjacency. In addition to validating the expected results of ligand-receptor pairs, identifying the proximity of potential source and destination cell groups responsible for that interaction can further influence the feasibility of that interaction **(Figure 5A)**.

**Figure 5:**
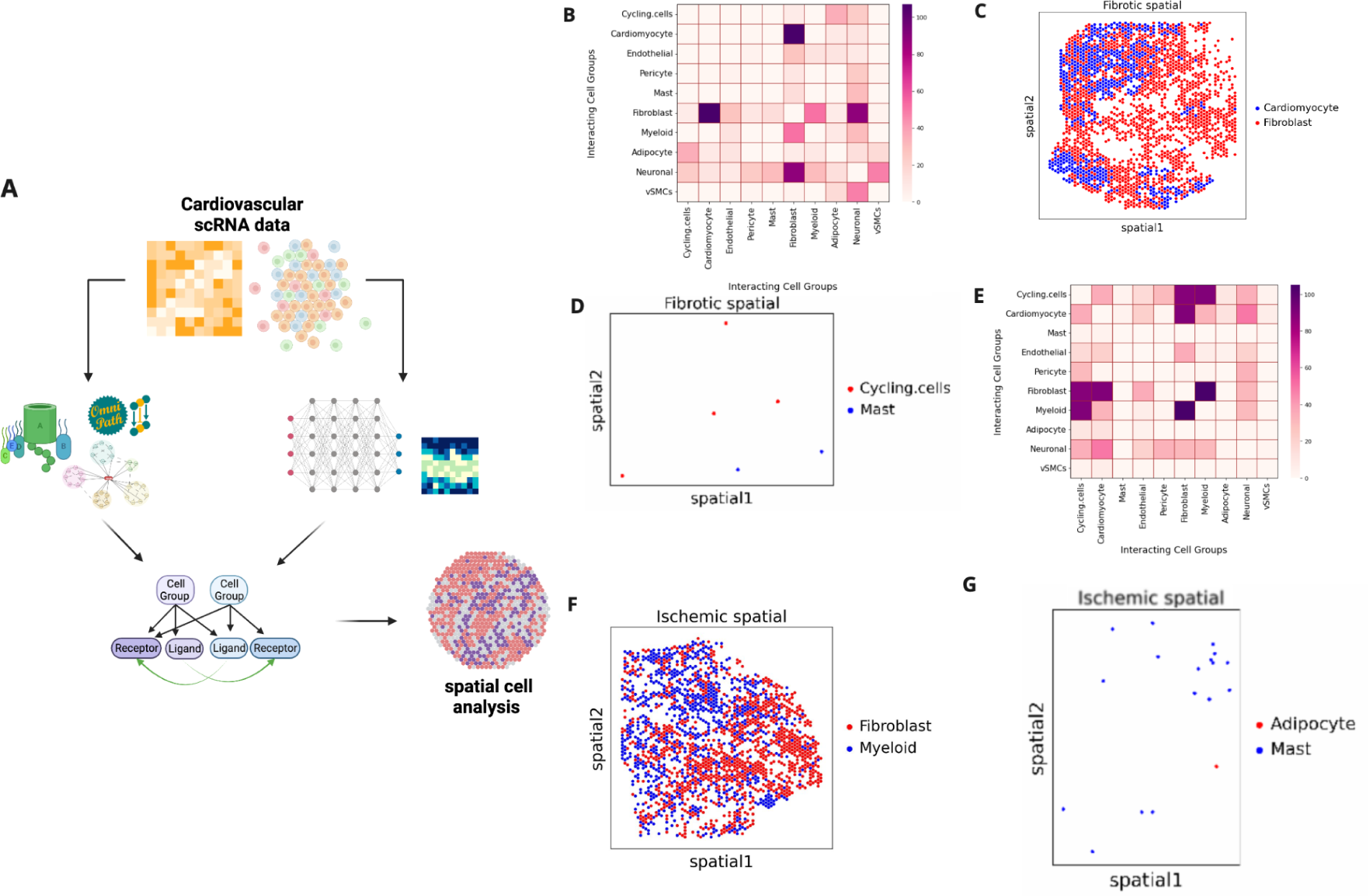
**(A)** GraphComm architecture can predict CCC occurring in spatially adjacent cell groups. Utilising spatial transcriptomics data from the fibrotic region of a patient sample, GraphComm can identify ligand-receptor interactions as well as identify potential source and destination cell groups. Then, the nodes of interest can be used to visualise two interaction cell groups and their respective proximity. **(B)** heatmap displaying number of interactions between any two cell groups, across all fibrotic slides **(C)** Spatial map only displaying cells in top interacting cell groups found by GraphComm in fibrotic slides, Cardiomyocyte and Fibroblast cells. From this spatial plot alone, the adjacency of these two cell groups and the potential for interactions can already be observed **(D)** Spatial map only displaying cells in low interacting cell groups found by GraphComm in fibrotic slides, Cycling cells and Mast cells. **(E)** Spatial heatmap displaying number of interactions between any two cell groups, across all ischemic slides **(F)** Spatial map only displaying cells in top interacting cell groups found by GraphComm in ischemic slides, Fibroblast and Myeloid cells. From this spatial plot alone, the adjacency of these two cell groups and the potential for interactions can already be observed **(G)** Spatial map only displaying cells in low interacting cell groups found by GraphComm in ischemic slides, Adipocyte cells and Mast cells. **Created in** https://BioRender.com

To assess how GraphComm can capture spatially probable interactions in different histomorphological regions of the heart, we performed CCC inference on 6 slides of the fibrotic zone and 8 slides of the ischemic zone from patients with myocardial infarction. The goal of this benchmark was to identify, across all slides available for a given zone, if GraphComm could consistently identify dominant patterns of spatially proximal and abundant cell groups. We found that in all fibrotic zone slides, GraphComm detected a large number of interactions between Cardiomyocyte cells and Fibroblast cells **(Figure 5B)**, whose spatial adjacency makes CCC highly probable **(Figure 5C)**. A more scarce interaction, between Cycling cells and Mast cells (**Figure 5D)** can be visualised as less spatially adjacent making the interaction less probable. Comparing these results quantitatively, these two cells groups’ euclidean distance can be measured to identify adjacency. Cell spatial adjacency is a value within the range of 0 and 1, with a lower spatial adjacency indicating more proximal cells overall and in this context more probable.Cardiomyocyte and Fibroblast cells are abundant across all fibrotic slides, with a mean euclidean distance of 0.02. Cycling cells and Mast cells are less abundant cell groups with a mean euclidean distance of 0.46.

Conversely, overall ischemic slides had a large number of interactions between Fibroblast and Myeloid cells **(Figure 5E)**, which is also probable as shown by spatial adjacency **(Figure 5F)**.Similarly to the fibrotic slides, less frequent interactions between Adipocyte and Mast cells **(Figure 5G)** is also less spatially adjacent. Myeloid and Fibroblast cells are abundant across all ischemic slides, with a mean euclidean distance of 0.019. Adipocyte and Mast cells have a more distant positioning with a mean euclidean distance of 0.068.

We then applied GraphComm on a single spatial transcriptomics dataset sample human heart obtained from the fibrotic zone (FZ) for CCC inference for a closer look at benchmarking GraphComm with spatial transcriptomics data, choosing the slide *FZ-GT-P19*. The dataset was accompanied by spatial coordinates for all cells, which could be used to construct a map of the tissue organisation by cell type (**Figure 6A)**. Based on GraphComm’s results, when analyzing only validated intercellular interactions, visual communication probability can be observed in dominant cell group interactions **(Figure 6B)**. To first identify what regions of the *FZ_GT_P19* slide could be indicative of potential intercellular activity, non-negative matrix factorization (NMF) was conducted on the slide using the framework LIANA+^33^ and visualised to determine the important cell groups for observing CCC **(Figure 6C)**. A top-ranked (rank 3) interaction found by GraphComm includes interactions between Myeloid cells and Neuronal cells, two cell groups with a mean spatial adjacency of 0.018 **(Figure 6D)**. This finding is also in accordance with important contributors to intercellular activity found with LIANA+, as the most highlighted slide region from the NMF analysis is a region containing Myeloid cells. In comparison, a low ranked cell group interaction is between Mast and vSMCs cells, whose interaction is less probable both in cell abundance and with a mean spatial adjacency of 0.13 (**Figure 6E).** In comparison to the NMF results as well, these cell groups are present in regions with lower value from the NMF. Against results from 100 randomization iterations the original prediction set of GraphComm had a lower median spatial adjacency of its 10 top ranked interacting cell types across all interactions. In comparison to the top 10 interacting cell types predicted by other CCC methods, GraphComm is able to outperform in predicting spatially closer cell groups, with the spatial adjacency of 4 interactions being lower in GraphComm’s prediction than all methods and 4 being lower in GraphComm’s prediction than 1 other method. **(Figure 6F).** These results show that with the correct information provided, GraphComm can not only prioritise biologically important and validated interactions but also prioritise interactions in more spatially adjacent cell types than other CCC methods.

**Figure 6:**
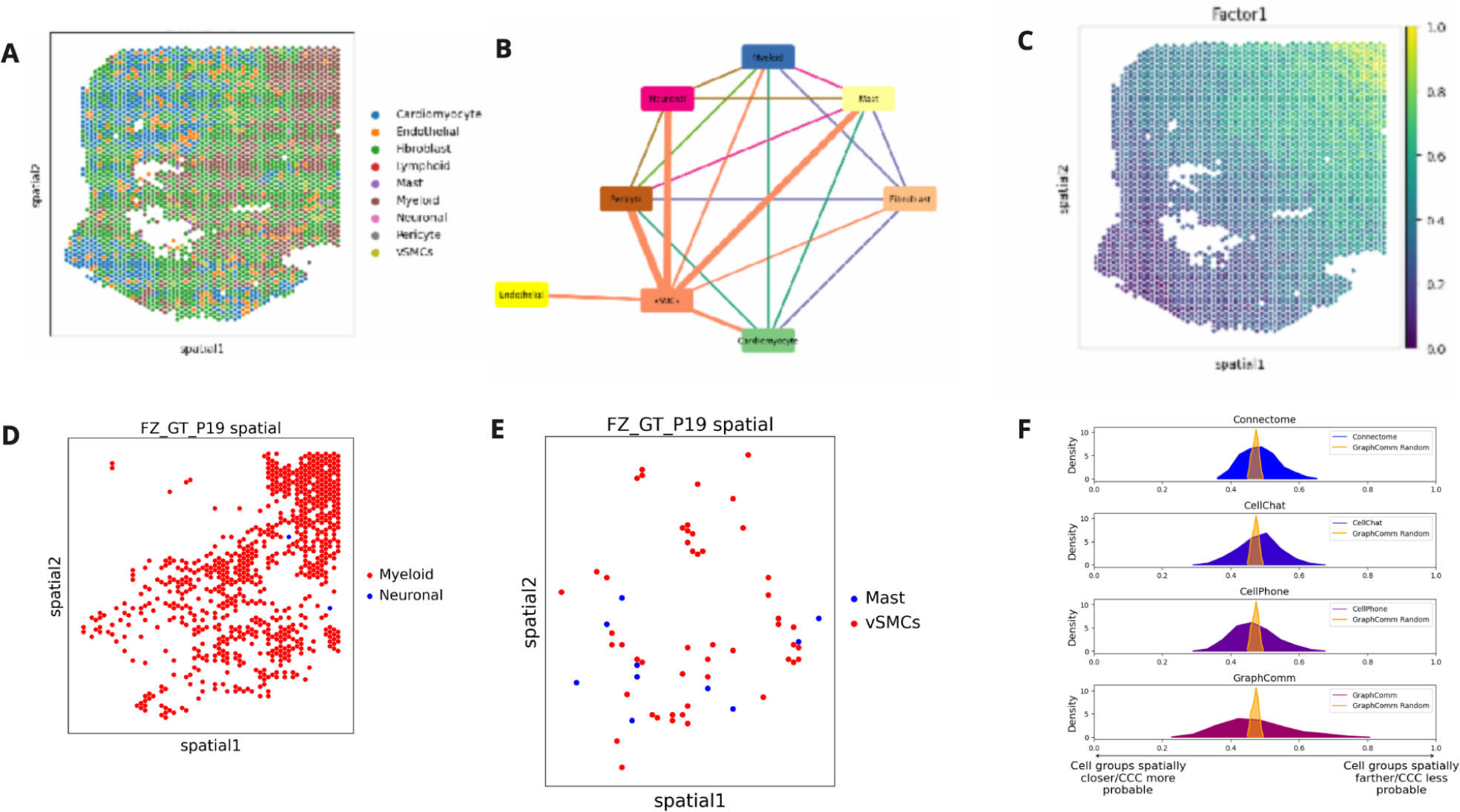
**(A)** Spatial coordinates of all cells in the dataset and colour coded by the cell-type in the *FZ_GT_P19* slide. **(B)** Weighted circle graph demonstrating the most prevalent cell group interactions found in the top 100 interactions predicted by GraphComm **(C)** Results from NMF done by LIANA+, detailing regions of the fibrotic slide most indicative of cell-cell communication **(D)** Spatial map only displaying cells in top interacting cell groups found by GraphComm, Neuronal and Myeloid cells. From this spatial plot alone, the adjacency of these two cell groups and the potential for interactions can already be observed **(E)** Spatial map only displaying cells in low-ranking interacting cell groups found by GraphComm, Mast and vSMCs cells. From this spatial plot alone, the adjacency of these two cell groups and the potential for interactions can already be observed **(F)** GraphComm had a lower median spatial adjacency of its 10 top ranked interacting cell types in comparison to results from 100 randomization iterations and other CCC methods.

## Discussion

The study of cell-cell communication has been found to provide novel insights into areas of biomedicine that were previously unexplored. By studying ligand-receptor interactions in cells of various cell types, development stage, disease condition or perturbation, critical processes involved in cell-state identity and function can be revealed. While many methods can predict possible CCC activity between cells, using a ground truth ligand-receptor database and transcriptomic data, these early methods have been found to not fully capture cell social networks as well as be limited in the incorporation of other modalities^5,11^. These initial results call for a need for a more comprehensive method for predicting CCC that captures the complex nature of cell-cell interaction networks, and includes prior ligand-receptor information in CCC prediction.

In this study, we introduce a new graph-based deep learning method, GraphComm, for inferring validated and novel cell-cell communication (CCC) ligand-receptor pairs in single-cell transcriptomic data. GraphComm is able to capture directed graphs demonstrating the relation between cell types and identified ligand/receptors as well as information for prioritising cell-cell interactions, and account for cell positioning, pathway annotation and protein complexes. We include both inter-cell and intracellular Protein-Protein Interactions (PPIs) obtained from OmniPath in our training phase. To account for CCC occurring both across and within one cell/cell type, GraphComm is trained on both intracellular and intracellular PPIs obtained from the OmniPath database so that GraphComm can be used for predicting CCC across and within one cell-type. For evaluation in the three use cases presented in this study, we validated using intracellular interactions for the mouse embryonic brain study so as to identify results that could reflect the original author’s outcome. For the other two datasets, we focus on intercellular communications to reflect the definition of cell-cell communication used in previous studies^11,17^ and focus on more diverse patterns that are occurring between different cell types/groups. We were able to prove that GraphComm can reproduce CCC results seen through spatial reconstruction, identify unique interactions in cell populations after drug treatment and prioritise interactions occurring in spatially adjacent cell types.

GraphComm’s capability to assign communication probabilities to ligand and receptor candidates enables various applications and innovations in future directions. Its successful performance in capturing diverse biological information from spatial cardiovascular datasets suggests potential applications in inference using multimodal data. In use cases such as predicting with gene expression coupled with other cellular data like ATACseq, GraphComm can provide a view of active CCC, as well as a more comprehensive analysis of cellular interactions and regulatory mechanisms. Moreover, GraphComm’s ability to uncover significant patterns in the PC9 cancer cell line dataset regarding chemoresistance and sensitivity pre- and post-treatment indicates its applicability to clinical datasets and different treatment conditions. This extends to scenarios involving recurrent cancers or instances where cells are predisposed to developing resistance over time. Lastly, GraphComm can be utilised in emerging technologies for representing single-cell data, such as simulated data.

Our study has several potential limitations. While GraphComm is able to produce some exciting results in identifying CCC interactions across multiple different datasets, there are still some unresolved challenges that exist in the field which require further investigation. In any given dataset, GraphComm identified on average 1000 source protein candidates and 1000 target protein candidates for interaction. Of all the possible interactions that could exist, a very small fraction of them were validated. Due to the nature of limited validated CCC activity, GraphComm and other methods in this field must combat a large class imbalance in learning social networks and filtering out the ground truth in a new dataset. Also, further exacerbated by the lack of ground truth, it has been previously recognized that there is a lack of gold standard for benchmarking in CCC with other methods. As research continues to develop and more validated CCC interactions are found in different contexts, predictions of ligand-receptor interactions and their evaluation will improve as well.

In conclusion, we anticipate that GraphComm provides a useful graph-based deep learning method that can accurately capture ligand-receptor events in single-cell transcriptomic data. The future application of GraphComm holds the potential to uncover valuable insights for a range of therapeutic and biomedical contexts, including the identification of key cell communication interactors that may serve as novel therapeutic targets in diseased cells.

## Methods

### Datasets

#### Single cell RNA datasets

GraphComm was benchmarked on three different datasets covering different biological settings - developmental, perturbation, and fibrosis/ischemia (**Table 1)**. All data used is available through public repositories (see Data Availability Section)

**Table 1:**
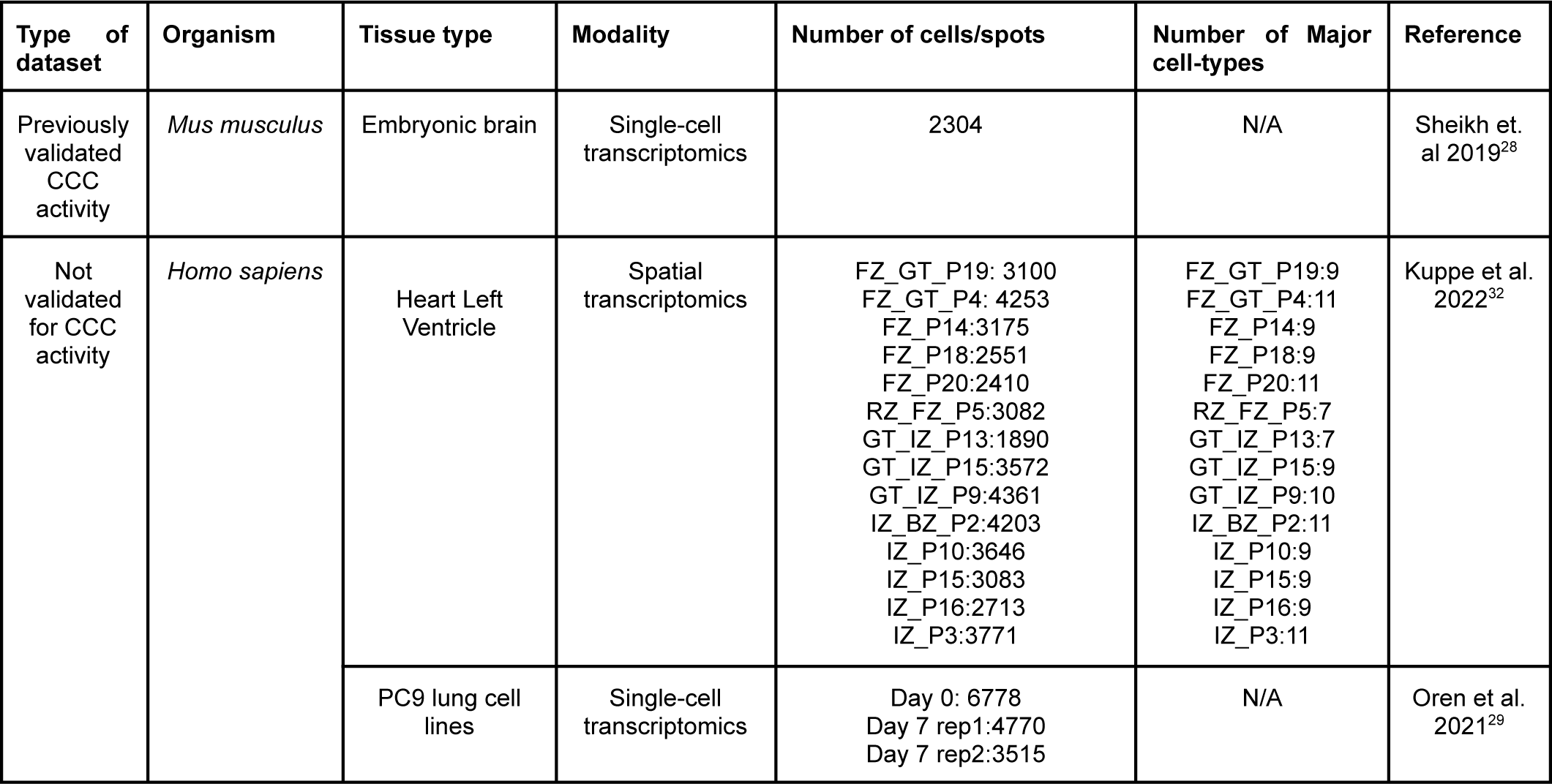
Single cell and spatial transcriptomics data used in benchmarking for GraphComm.

| Type of dataset | Organism | Tissue type | Modality | Number of cells/spots | Number of Major cell-types | Reference |
| --- | --- | --- | --- | --- | --- | --- |
| Previously validated CCC activity | *Mus musculus* | Embryonic brain | Single-cell transcriptomics | 2304 | N/A | Sheikh et. al 2019<sup>28</sup> |
| Not validated for CCC activity | *Homo sapiens* | Heart Left Ventricle | Spatial transcriptomics | FZ_GT_P19: 3100<br>FZ_GT_P4: 4253<br>FZ_P14:3175<br>FZ_P18:2551<br>FZ_P20:2410<br>RZ_FZ_P5:3082<br>GT_IZ_P13:1890<br>GT_IZ_P15:3572<br>GT_IZ_P9:4361<br>IZ_BZ_P2:4203<br>IZ_P10:3646<br>IZ_P15:3083<br>IZ_P16:2713<br>IZ_P3:3771 | FZ_GT_P19:9<br>FZ_GT_P4:11<br>FZ_P14:9<br>FZ_P18:9<br>FZ_P20:11<br>RZ_FZ_P5:7<br>GT_IZ_P13:7<br>GT_IZ_P15:9<br>GT_IZ_P9:10<br>IZ_BZ_P2:11<br>IZ_P10:9<br>IZ_P15:9<br>IZ_P16:9<br>IZ_P3:11 | Kuppe et al. 2022<sup>32</sup> |
|  |  | PC9 lung cell lines | Single-cell transcriptomics | Day 0: 6778<br>Day 7 rep1:4770<br>Day 7 rep2:3515 | N/A | Oren et al. 2021<sup>29</sup> |

#### Curated ligand-receptor database

In selecting a database that could be used to establish a ground truth for cell-cell communication, we have decided to source all validated ligand-receptor interactions from the OmniPath database. Specifically, all scientifically validated PPIs (30,053 pairs), intercellular interactions (3,731 pairs) protein complex grouping (8,022 complexes), and KEGG pathway annotation (7,534 entries) were extracted from the OmniPath database via the python package *omnipath* 1.0.6^22,34^. Data extracted using this python package was provided in a series of tables, using HGNC symbols for intracellular proteins/ligands/receptors as identifiers.

### Preprocessing of single-cell data and creation of a directed cell to LR graph

To prepare single-cell RNAseq data into a computable input for predicting cell-cell communication, raw counts were converted to a log-transformed normalised count. Significantly expressed genes were identified for each cell group via identifying genes that had an average non-negative expression across all cells of a given cell type. For each cell group, the set of genes was subsetted to strictly include HGNC identifiers that also appear in the OmniPath database as a potential ligand/receptor or intracellular protein. Then a directed graph was constructed (using Pytorch Geometric^35^) pointing from each cell group to their respective significant intracellular protein/ligand/receptor.

### Model training and inference

CCC predictions via GraphComm are generated via a two step pipeline: first building a prior model using the OmniPath Database, and then performing inference using transcriptomics data.

#### Building of prior model using OmniPath database

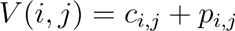

After identifying ligands and receptors present in the scRNA dataset based on their expression profile, the OmniPath database is cross referenced to identify validated ligand-receptor present within the set. A directed graph is constructed using the Pytorch Geometric^35^ library with each protein as a node, and validated edges are drawn if a pair of proteins participate in a validated intracellular or intercellular interaction. Alongside the directed graph, a matrix is constructed with the dimensions (# of nodes) x (# of nodes). For each cell in the matrix, which points to the intersection of any two nodes, the value is as follows:

Where *V(i,j)* is the value for that cell in the feature matrix, *c_i,j_* is the number of protein complexes that node *i* and *j* are both subunits in and *p_i,j_* are the number of KEGG pathways that *i* and *j* are both members of. The constructed directed graph is then trained for 100 epochs through Random Walk via a Node2Vec^25^ Model, with an output embedding dimension of 2. The output after the training period is two dimensional positional information for each node in the input graph. Inner products of output node embeddings are used to construct a positional matrix with the shape of number of source proteins x number of target proteins and then multiplied with the constructed annotation matrix to further scale the output in accordance with protein complex and pathway information.

### Inference on cell-cell communication using transcriptomic data and Graph Attention Network

#### Construction of Cell and Ligand/Receptor embedding matrix

Determining the ranking of ligand-receptor pairs begins with the directed graph pointing cell groups to ligands and receptors, as constructed in the preprocessing step. Similar to the graph used in building a prior model, before inference, a feature matrix of shape *(# of nodes) x (# of nodes)* is constructed to further annotate the directed graph. These can be recognized as or referred to as biological node embeddings, as they capture dataset specific information from a biological context.

The value for each cell, which points to the intersection of any two nodes, are as follows for node *i* and node *j:*

If node *i* is a source/target protein and/or node *j* is a source/target protein:

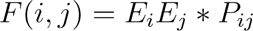

Where *E_i_*is the average expression of source/target protein *i, E_j_* is the average expression of source/target protein *j* and *P_i,j_*is the value corresponding to the matrix generated by the previous prior model step.

If nodes *i* is a cell group and node *j* is a source/target protein:

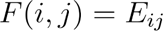

Where *E_ij_*is the average expression of source/target protein *j* across cells of cell group *i*.

All numerical values regarding gene expression in the node embedding matrix are scaled to be in the range between 0 and 1, to match the scale of cell group information.

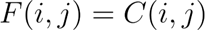

If nodes *i* and *j* are both cell types, embedding is only provided in the event that there is cell-specific information accompanying the dataset that is indicative of positional information. An example of this could be spatial coordinates for cells.In that case, the embedding value is the following:

Where *C(i,j)* is a function of choosing that resembles the relation between cell type *i* and *j* from positional information. For spatial coordinates, for example, *C(i,j)* is computed as the minimum euclidean distance between any cell of type *i* and cell of type *j*.

### Model Architecture

We then passed the directed graph from the preprocessing step, with the corresponding embedding matrix, through a Graph Attention Network (GAT).^27^ GATs use attention mechanisms to update pre-defined information about nodes depending on their connectivity. Any given attention mechanisms *a* is computed using the following equation from the original publication^27^:

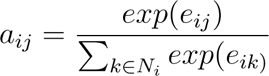

where i is a given cell/ligand/receptor node, *j/k* are nodes in the direct neighbourhood of node i, *N_i_* is the total set of nodes and *e_ij_*/e_ik_ are coefficients computed using input node embeddings and trainable weight matrices for each node. Attention mechanisms are then summed for all neighbouring nodes to obtain an updated node embedding for node *i* **(Extended Data Figure 2)**. Attention mechanisms are updated for 100 epochs during the training process. After being passed through a log softmax function, node embeddings for ligand and receptor nodes are utilised to minimise cross entropy loss^36^ against assigned class labels for input ligands and receptors (1 if a ligand/receptor participates in a validated intercell interaction, 0 if not). The loss function can be described as the following (Equation 2):

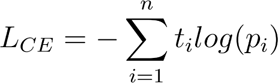

Where *L_CE_*is the cross-entropy loss, *N* is the number of nodes, *t_i_*is the truth label (in this case, 1 or 0) and *p_i_* is the sigmoid probability for that node from GraphComm.

### Inference using trained GAT model

A Graph Attention Network was chosen for the prediction step of ligand-receptor interactions due to its learning from an embedding matrix of values across a continuous range, which could contain information for different node types, and its design for node classification. After the training period, a single forward pass is done to receive a two-dimensional numerical array containing updated embeddings for all nodes in the graph. The dimensions of the updated embeddings correspond to, for each node, the probability of it being labelled as class 0 (non interacting) or class 1 (interacting). To obtain top ranked validated LR links, an inner product is done of positive class probabilities for all ligand and receptor candidates (indicating the probability that they are participating in an interaction). The output is a matrix of the shape *(# of ligands) x (# of receptors).* This matrix is then flattened to a shape of *(# of possible protein pairs) x 1* and ranking of LR links is decided by sorting the inner products in descending order.

To obtain source cell and destination cell types, an inner product is computed of the ligand and receptor embeddings by the cell type embeddings from the first output of the Graph Attention Network. This returns a matrix of the shape *(# of ligands)* and *(# of receptors)*, assigning communication probability for every possible edge. Interactions are ranked by inner product value.

### Randomisation Experiments

The randomization experiments on a given scRNA dataset are done via performing the same method of communication modelling, with the same values for annotations for graphs, but by randomly shuffling the edges of both the OmniPath and transcriptomic input graph. This will cause the model to perform inference using a randomised ground truth and with different assignments of cell groups to ligands and receptors. For each dataset, randomization was done for 100 iterations.

## Data Availability

The sequenced embryonic mouse brain cells and PC9 cell lines used in this study are publicly available in Gene Expression Omnibus (GEO) at GSE133079^28^ and GSE150949^29^. Spatial transcriptomics data^32^ used in this study is publicly available via the DOI 10.5281/zenodo.6578047.

## Code availability

The code for GraphComm and notebooks used to analyse data presented in this study are provided in the GitHub repository https://github.com/bhklab/GraphComm and in a Code Ocean capsule.^37^

## Research Reproducibility

Full reproducibility of all results is available at the corresponding <u>Code Ocean Capsule</u>^37^. Model checkpoints, as well as results used to generate published figures are available for analysis. The Code Ocean capsule is configured to, at the click of a button, run pre-saved models on all inference datasets included in this manuscript.

## Acknowledgements

We thank D. Forster, H. Cui, H. Maan and I. Smith for their insightful comments. Infographics in figures were created with Biorender. Experiments were run using resources from the Vector Institute.

---

**Extended Data Figure 1:**
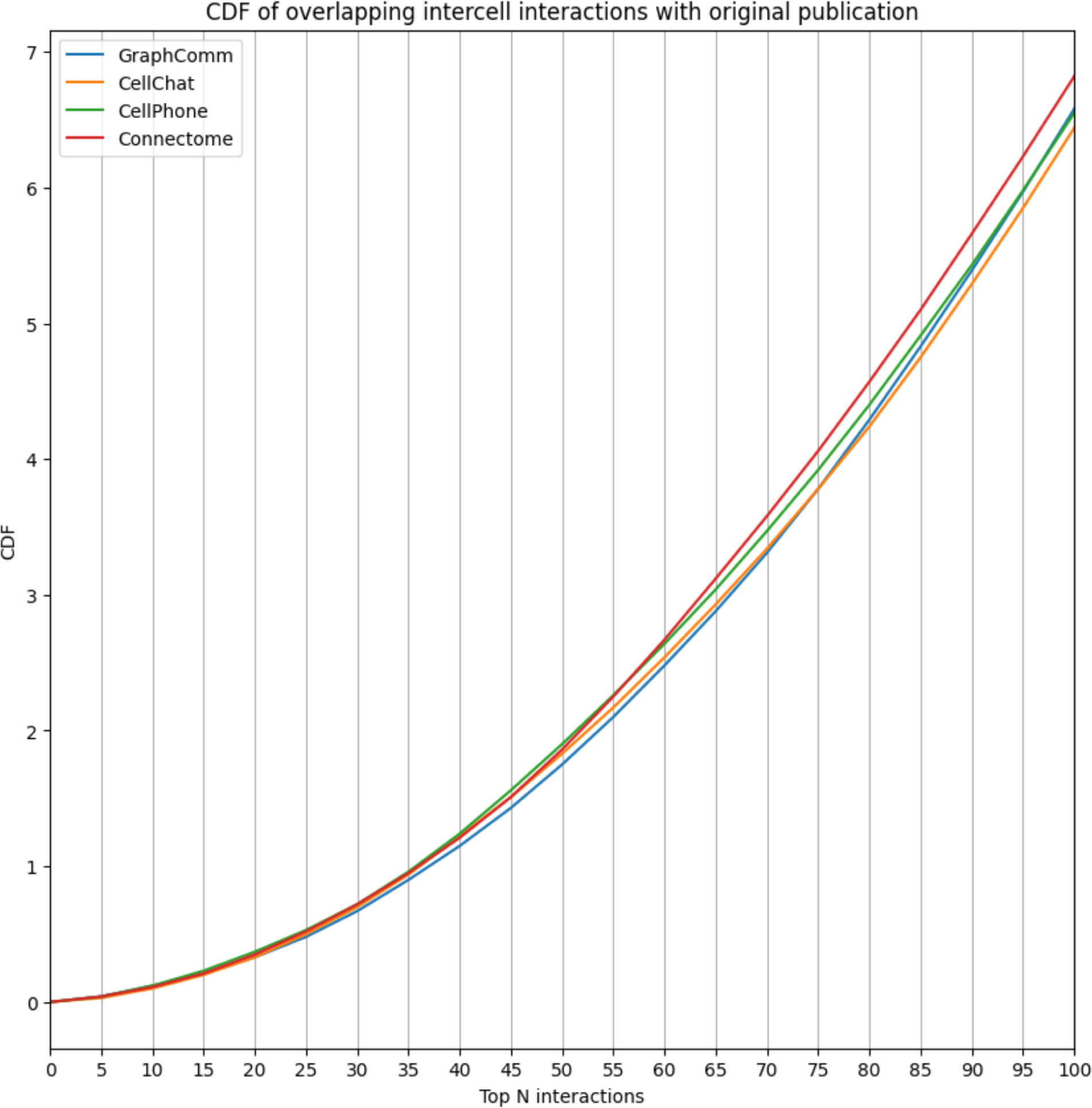
In the top 100 interactions, GraphCom predicts a comparable or slightly lower number of interactions that overlap with the publication in comparison to other methods. Potential reasoning behind this result includes GraphComm’s prioritisation of previous intercellular interactions that were not originally investigated.

**Extended Data Figure 2:**
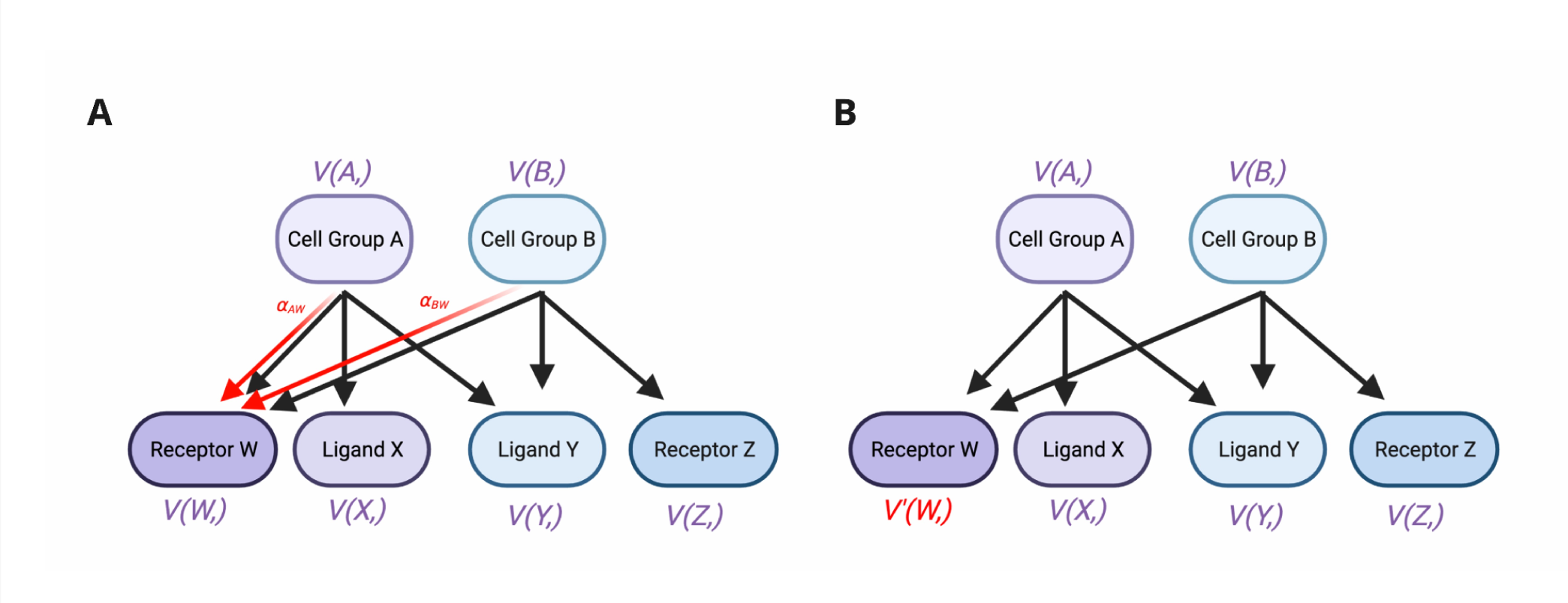
The mechanisms behind the GAT architecture, demonstrating how embedding information is updated for nodes during model training. **(A)** Input graph during GAT training, during embedding update of a single node, Receptor W (all nodes are updated during one training epoch). A given node *N* is annotated with V(*N,)*, representing the full vector of embeddings (Node *N’s* numerical relation to all other nodes in the graph). From each first order neighbouring node of Receptor W, the attention mechanism with respect to Receptor W is calculated and concatenated to the existing Receptor W embedding. **(B)** Input graph post update of Receptor W embedding, *V(W,)* has been updated to *V’(W,)* to reflect the updated embedding. The process is continued for each node in the graph, which completes the epoch. Both target nodes are updated from their source nodes and vice versa (i.e. embeddings from ligands and receptors are used to update cell group embeddings). **Created in** https://BioRender.com

